# Divergent excitation two photon microscopy for 3D random access mesoscale imaging at single cell resolution

**DOI:** 10.1101/821405

**Authors:** FK Janiak, P Bartel, MR Bale, T Yoshimatsu, E Komulainen, M Zhou, K Staras, LL Prieto-Godino, T Euler, M Maravall, T Baden

**Affiliations:** Sussex Neuroscience, School of Life Sciences, University of Sussex, UK; The Francis Crick Institute, London, UK; Institute of Ophthalmic Research, University of Tuebingen, Germany; Centre for Integrative Neuroscience, University of Tuebingen, Germany

## Abstract

In neuroscience, diffraction limited two-photon (2P) microscopy is a cornerstone technique that permits minimally invasive optical monitoring of neuronal activity. However, most conventional 2P microscopes impose significant constraints on the size of the imaging field-of-view and the specific shape of the effective excitation volume, thus limiting the scope of biological questions that can be addressed and the information obtainable. Here, employing ‘divergent beam optics’ (DBO), we present an ultra-low-cost, easily implemented and flexible solution to address these limitations, offering a several-fold expanded three-dimensional field of view that also maintains single-cell resolution. We show that this implementation increases both the space-bandwidth product and effective excitation power, and allows for straight-forward tailoring of the point-spread-function. Moreover, rapid laser-focus control via an electrically tunable lens now allows near-simultaneous imaging of remote regions separated in three dimensions and permits the bending of imaging planes to follow natural curvatures in biological structures. Crucially, our core design is readily implemented (and reversed) within a matter of hours, and fully compatible with a wide range of existing 2P customizations, making it highly suitable as a base platform for further development. We demonstrate the application of our system for imaging neuronal activity in a variety of examples in mice, zebrafish and fruit flies.

---

Laser scanning two photon (2P) microscopy allows the imaging of live cellular processes deep inside intact tissue with high signal-to-noise, temporal fidelity and spatial resolution (Denk et al. 1990). Nonetheless, standard diffraction-limited 2P setups with a collimated laser excitation beam have several key characteristics that constrain their broad applicability; namely, a typically small field of view (FOV), a fixed-size excitation spot and restricted options for rapid random access 3-dimensional scans. These are significant limitations because the biological samples that are interrogated with 2P microscopy can exhibit substantial variations in size and spatial structure. For example, the volume of an adult mouse brain is approximately four orders of magnitude larger than that of a larval zebrafish, and seven orders of magnitude larger than a first instar larval fruit fly (Fig. 1a). Similarly, neuronal sub-structures are also highly variable in size, ranging from sub-micron levels for synapses to >20 μm for somata. Additionally, neural densities vary by more than an order of magnitude across different animal brains (Weisenburger & Vaziri 2018). As such, 2P microscopy tends to reveal very different levels of detail and organization across its diverse experimental applications. To maximize biological information, upgrades for 2P microscopy should enable the imaging of neuronal activity from many neural structures of a given size and density across a sufficiently large 3D volume of tissue at sufficiently high frame rates for the chosen neuronal process and biosensor while maintaining single-cell or-synapse resolution.

**Figure 1.**
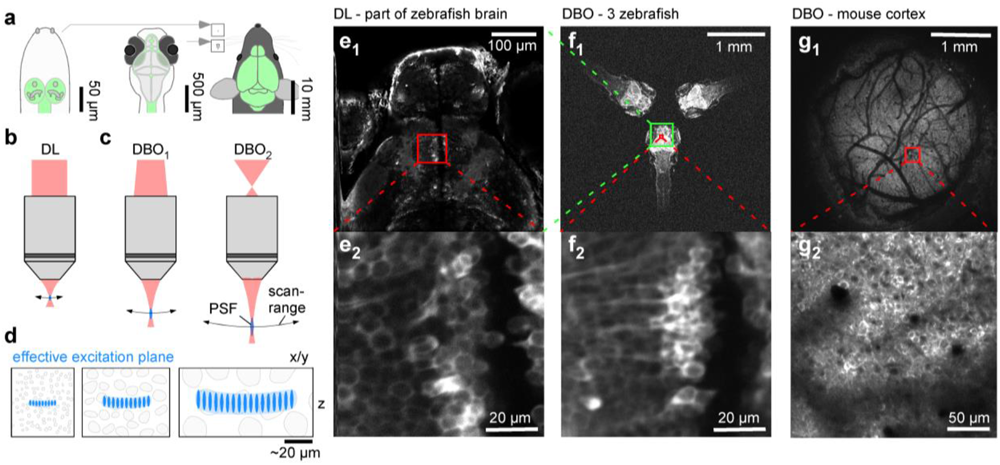
Field of view expansion through divergent beam optics. **a**, Schematics of larval Drosophila (left), larval zebrafish (centre) and adult mouse (right) with central nervous system highlighted (green) to illustrate size differences. Insets next to the mouse for direct size-comparison between these species on the same scale. **b**, optical configurations standard diffraction limited (DL) 2P setup with parallel laser beam entering objective’s back aperture. **c**, divergent beam optics (DBO) configurations use a still diverging laser beam instead. As a result, the field of view and focal distance are expanded and the point spread function (PSF) becomes elongated. These effects scale with the angle of divergence: left, mild divergence (DBO_1_), right, strong divergence (DBO_2_). **d**, Schematic representations of typical neuronal somata in species shown in (a), as interrogated by 2P setups shown in (b,c), respectively. **e**, in vivo 7 dpf larval zebrafish (HuC::GCaMP6f) imaged with an out-of-the-box Sutter MOM DL setup at full field of view (e_1_) and when zoomed in to reveal individual neuronal somata (e_2_) as indicated. **f**, same zebrafish as shown in (e), as well as two further zebrafish imaged using our DBO configuration at maximal field of view. **g**, In vivo adult mouse cranial window over somatosensory cortex imaged with DBO maximal field of view (g_1_) and when zoomed in as indicated (g_2_).

In response to this demand, a profusion of custom modifications to 2P microscopes have been developed to expand the spatial and temporal boundaries over which neural structures can be optically interrogated. For example, the maximal planar field of view (FOV) has been increased from typically 0.5 mm to between 3.1-10 mm diameter by the exchange of optical components (7 mm) (Bumstead 2018), large diameter off-the-shelf optical components (10 mm) (Tsai et al. 2015), custom built objectives (3.1 mm) (Stirman et al. 2016) and a mesoscope configuration (5 mm) (Sofroniew et al. 2016) to allow ‘mesoscale’ interrogation of neural circuits. In parallel, customizations using multiple beams have allowed simultaneous scanning of distant brain regions (Cheng et al. 2011; Han et al. 2019; Stirman et al. 2016). Likewise, higher temporal resolutions have been achieved by tailoring the point spread function (PSF) to the geometry and distribution of the neuronal structures of interest, thus increasing signal-to-noise ratio (SNR) and, in turn, decreasing the minimally-required dwell time per pixel (Prevedel et al. 2016; Weisenburger et al. 2019). Moreover, the imaging plane has been axially expanded by engineering an excitation spot with Bessel focus (Botcherby et al. 2006; Lu et al. 2017) or by elongated Gaussian foci stereoscopy (Song et al. 2017). These customizations provide efficient ways to merge image structures that are located at different depths into a single volumetric plane. Furthermore, in recent years, systems integrating acousto-optic deflectors (Chong et al. 2019; Grewe et al. 2010), electrical tunable lenses (Grewe et al. 2011; Han et al. 2019; Sheffield & Dombeck 2015; Yang et al. 2018; Zhao et al. 2019) and remote focusing units (Botcherby et al. 2006; Sofroniew et al. 2016) have enabled quasi-simultaneous multiplane volumetric scans.

These types of extensions have been essential in driving the field forward, yet many are expensive, require custom-produced optical elements, complex optical alignment and/or introduce new limitations. The latter can include limitations in both excitation (e.g. power loss (Sofroniew et al. 2016), waterfront dispersion (Chong et al. 2019)) and collection (Bumstead 2018; Tsai et al. 2015). Here we introduce a new optical design for 2P microscopy that overcomes many of these limitations while simultaneously matching the capabilities of a wide range of state-of-the-art performance customisations, while being ultra-low-cost, simple and flexible.

Implemented for ~£1,000 on an existing 2P setup equipped with a standard x20 objective, our “divergent beam optics” (DBO) design allows the expansion of the planar FOV from typically ~0.5 mm in diameter to anywhere up to 3.5 mm to flexibly suit experimental needs (Fig. 1). This expansion is accompanied by only a moderate and adjustable increase in the system’s 3D PSF while maintaining single cell resolution over a wide range of biological applications. Indeed, our approach affords the possibility to flexibly tune the PSF’s 3D shape to the experimental needs (“PSF tailoring” (Weisenburger et al. 2019)). Overall, the DBO design increases a standard diffraction limited (DL) microscope’s space bandwidth product (i.e. ~the total amount of information, or number of pixels, the microscope can acquire in a single frame; (Lohmann et al. 1996)). Effectively, what is lost in spatial resolution due to *PSF* expansion is more than made up for by the larger FOV. For example, unlike a standard diffraction limited (DL) setup (Fig. 1b,e), our DBO setup (right in Fig. 1c,f_1_,g_1_) allows simultaneous imaging of three entire zebrafish brains, or about a third of the width of a mouse’s cortex, while in each case maintaining single cell resolution (Fig. 1f_2_,g_2_, Supplementary Video S1).

Another key feature of our design is that it allows near-simultaneous sampling in distant brain regions separated in 3 dimensions. This is achieved by the introduction of an electrically tunable lens (ETL) into our DBO setup permitting up to 0.6 mm axial-jumps within a single scan-line and thus providing millisecond-precision arbitrary guidance of the laser-focus between any two points within a 3.5 (diameter) x 0.6 mm (height) cylinder. We also demonstrate how this introduces the possibility to “bend” an imaging plane in three dimensions to better account for natural curvature in biological samples. The setup is controlled by the free and open-source custom 2P scan software ScanM (by T Euler and M Mueller) as well as a standalone graphical user interface (GUI) that communicates with microcontrollers to control the ETL, Pockels cell and light emitting diodes (LEDs) in synchrony with the mirror commands (Franke et al. 2019) e.g. for visual stimulation or optogenetics. This independence of microcontroller controls from the scan software means that our system can be seamlessly integrated with any existing software solution (e.g. ScanImage (Pologruto et al. 2003)).

In summary, our DBO design offers a large volumetric field of view for rapid random access multiplane imaging or the execution of arbitrary 3D scan paths. Crucially, our solution is both comparatively low-cost and easy to implement on any existing 2P setup without the need for complex optical calibration, thus facilitating its widespread adoption in the community. We anticipate that others will be able to build on our core optical design using existing and new modifications to further increase its capability in the future. We demonstrate the current performance of our system with a range of examples from mice, zebrafish and fruit flies.

## RESULTS

### Divergent beam optics for field of view expansion

In traditional laser scanning 2P microscopy (Fig. 1b), a diffraction limited (DL) *PSF* is generated to excite fluorophores in a typically sub-micron volume of tissue. Here, xy-scanning mirrors reflect the laser beam into a collimation system comprised of a scan and a tube lens. The collimated beam then enters the back aperture of a high numerical aperture (N.A.) objective (Denk & Svoboda 1997; Svoboda & Yasuda 2006) to converge the parallel rays into a DL point at focal distance (Born & Wolf 1980). The Gaussian shape of the excitation beam dictates that it is not possible to perfectly match beam width to the objective’s back aperture. Instead, the back aperture is typically overfilled with a factor of 1/e^2^ as a compromise between maximising spatial resolution (i.e. small *PSF* size) and power transmission (Philbert S. Tsai and David Kleinfeld 2009).

In contrast, our DBO design (Fig 1c) illuminates the objective’s back-aperture with a decollimated and divergent beam (hence “divergent beam optics”, DBO). This leads to an increased angle of view as the light exits the objective mouth, such that the same angular scan-mirror movement leads to a larger absolute shift in the image plane – thereby greatly increasing the FOV. In parallel, this also alters the effective excitation numerical aperture (N.A.) to yield a larger-than-DL excitation spot (i.e. an elongated *PSF*) at greater focal distance. The magnitude of each of these effects, scales with the angle of divergence as the beam enters the back aperture of the objective. Accordingly, simply shifting the planoconvex lenses up or down the laser path, or switching between different refractive power lenses, provides for easy control over the system’s optical properties to flexibly suit the user’s needs.

In the following we show that the use of DBO in 2P microscopy brings about important advantages over the traditional, collimated and diffraction limited (DL) design:

1. The total field of view (FOV) can be expanded several-fold to suit the user’s needs.
2. Scan-mirror movements translate into correspondingly larger xy-shifts in the image plane, meaning that even multi-millimetre random access jumps can be achieved with millisecond precision.
3. The addition of an electrically tunable lens (ETL) in front of the scan mirrors allows for similarly extensive expansion in the axial dimension.
4. The simplified optical path and under-filling of the objective’s back aperture means more laser power is available at the sample plane.
5. It allows flexible and partially FOV-independent *PSF*-shape adjustment for imaging neurons of different size to individually optimise detection sensitivity for different biological samples (Prevedel et al. 2016; Weisenburger & Vaziri 2018).
6. It increases the space-bandwidth product (SBP) (Bumstead 2018; Lohmann et al. 1996), meaning that more information can be transmitted for a given FOV.
7. The increased working distance provides additional space for access to the preparation, for example with electrodes or stimulation equipment.

We first discuss the required optical modifications and their impact on key excitation parameters (Fig. 2, Figs. S1–3), before presenting a series of key use cases of different configurations for the interrogation of neural structure and function across diverse models (Fig. 3–7, Fig. S4).

**Figure 2.**
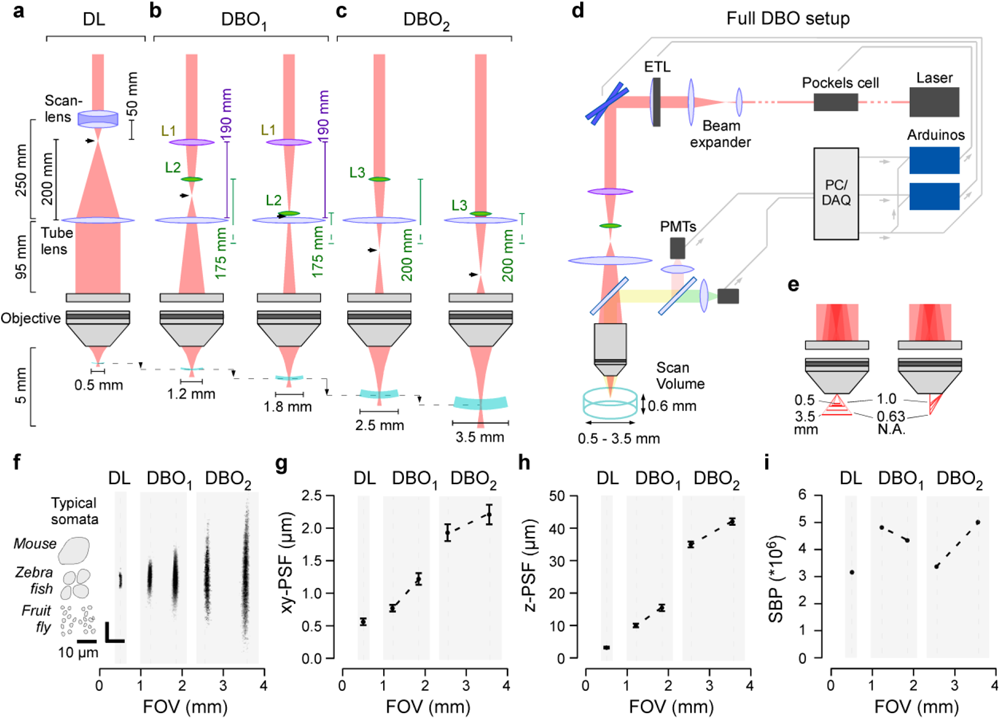
I Divergent beam optics in 2P microscopy. **a**, Optical configuration of a standard DL setup with collimation system consisting of a scan lens and a tube lens to set-up an infinity collimated laser beam at the level of the objective’s back aperture. Effective refractive power and relative distances of lenses indicated. The intermediary focal point (IFP) is immediately behind the scan lens (arrowhead). **b,** DBO_1_ configuration replaces the scan lens with a pair of plano-convex lenses (L1,2) as indicated. The relative position of L2 to the tube lens defines the position of the new IFP, which is now further along the laser path. As a result, the field of view can be expanded to between 1.2 and 1.8 mm. **c**, DBO_2_ configuration as (b), but now using a single plano-convex lens (L3) allows FOV expansion to 2.5 – 3.5 mm. **d**, complete DBO setup, including an ETL positioned in front of the scan mirrors for rapid z-scanning. PMTs, Photomultipliers. **e**, FOV expansion under DBO combines two effects: Increased focal distance (left) and reduced numerical aperture (N.A., right), which together give rise to a larger effective focal plane and enlarged PSF. **f**, point spread functions (PSFs) measured for all optical configurations, with size of typical neuronal somata of different species indicated. **g,h**, lateral (g) and axial (h) spread of the PSFs quantified. Errors in s.d. **I**, space-bandwidth product (SBP) of the different configurations.

**Figure 3.**
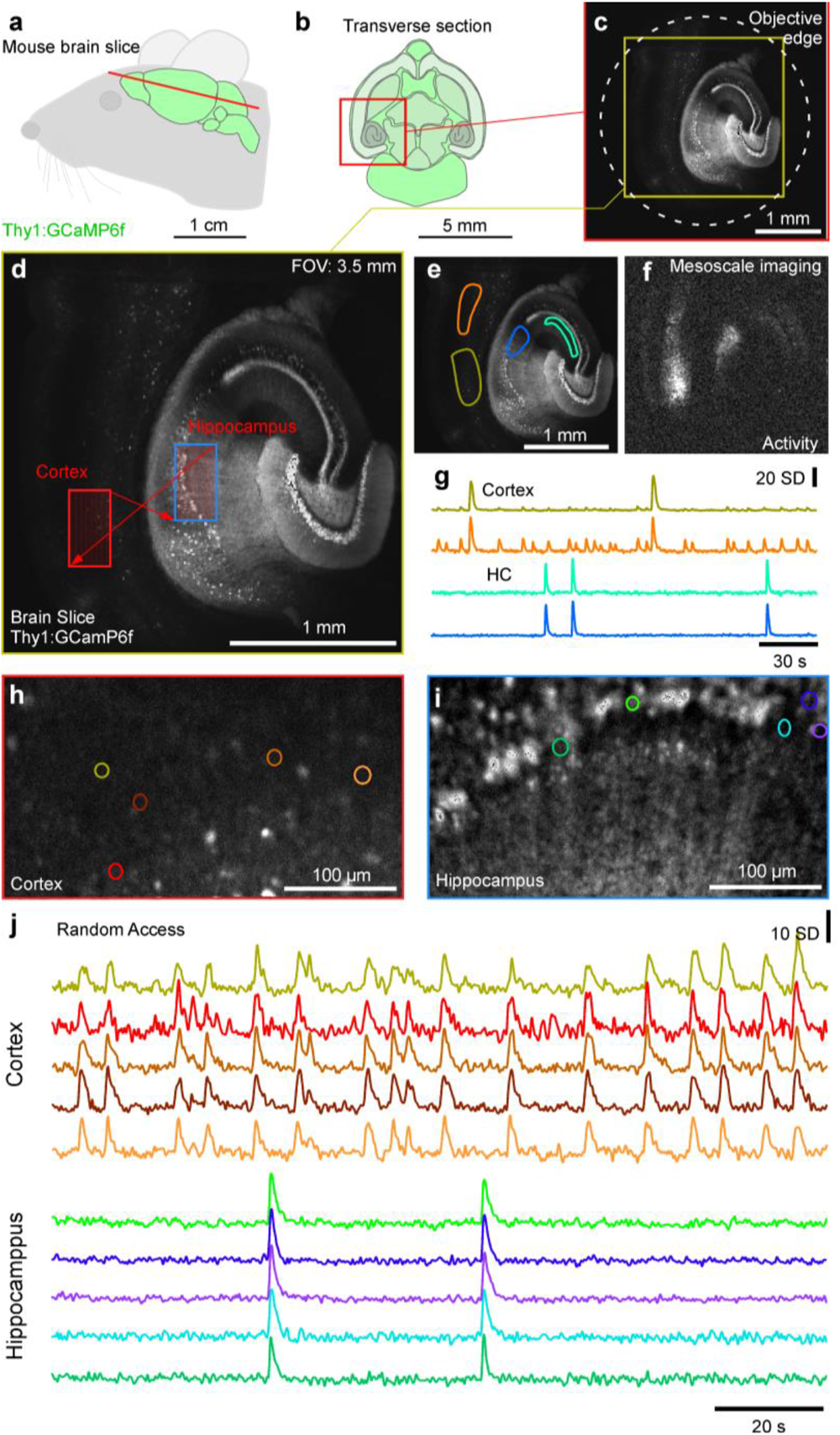
High-resolution mesoscale and random access imaging: brain slice. **a**,**b**, Schematic of brain (a) and transverse section (b) of a Thy1:GCaMP6f mouse. **c**,**d**, 1024×1024 px DBO_2_ example scan of slice through cortex and hippocampus at maximal FOV (c) and DBO_2_ zoom in (d) as indicated. The slice was bathed in an epileptogenic (high K^+^, zero Mg^2+^) solution to elicit seizures. **e-g**, Mean of 256×256 px scan (3.91 Hz) of (d) with regions of interest (ROIs) indicated (e), activitycorrelation projection (Methods) indicating regions within the scan showing regions of activity (f) and z-normalised fluorescence traces (g). **h-j**, 2 times 128×256 px (3.91 Hz) random access scan of two regions as indicated in (d) allows quasi-simultaneous imaging of the cortex (h) and hippocampus (i) at increased spatial resolution, with fluorescence traces (j) extracted as in (d,e).

## PART I. OPTICS, EXCITATION AND OPTICAL SAMPLING

### A simple scan-lens modification yields up to 7-fold FOV expansion

An off-the-shelf infinity-corrected galvo-galvo Sutter-MOM setup equipped with a 20x objective (Zeiss Objective W “Plan-Apochromat” 20×/1.0) offers a FOV diameter of ~0.5 mm (Fig. 2a). However, when underfilling the back aperture of the objective with a diverging laser (Fig. 2b-d), the beam exits the objective front aperture at increasingly obtuse angles at an effectively decreased N.A. and comes into focus at a greater distance (Fig 2e, Fig. S1a). Together, this expands the effective excitation FOV in both xy (increased angle and decreased N.A.) and z (elongated PSF). To achieve this effect, it is necessary to bring the collimated laser beam, having passed the scan mirrors, to an “early” intermediary focal point (IFP) prior to reaching the objective, thus setting up the diverging beam thereafter (Fig. 2b,c, arrowheads). The exact divergence angle as the beam enters the back-aperture of the objective, and thus the magnitude of the abovementioned effects, then depends on the position of this IFP in the laser path. We present two simple optical solutions (DBO_1_ and DBO_2_) to set-up an early IFP and thus expand the effective FOV to varying degrees.

In the standard DL configuration, the scan-lens (SL) and tube lens (TL) are separated from each other at a distance that is equal to their combined focal lengths (50_SL_ + 200_TL_ mm = 250 mm) to collimate the beam (Fig. 2a). In DBO_1_, we removed SL and instead inserted two off-the-shelf plano-convex lenses (L1, modified VISIR 1534SPR136, Leica; L2, LA1229 Thorlabs) with focal lengths 190 and 175 mm, respectively (Fig. 2b, Methods). L1 was fixed 190 mm in front of TL to set up an IFP exactly at the TL. Next, L2 was positioned between L1 and TL to further increase laser convergence and thus shift the exact position of IFP away from the TL. Accordingly, IFP is always in front of the TL, with L2 determining its exact position: Simply shifting L2 along the laser path between 100 and 5 mm distance from the TL expanded the effective FOV diameter to anywhere between 1.2 and 1.8 mm, respectively (compare Fig. 2b left and right). In DBO_2_ (Fig. 2c), we replaced SL with a single lens (L3) of 200 mm focal length (LA1708, Thorlabs). L3 operated in much the same way as L2 in the previous modification M1, however now the IFP was behind rather than in front of TL. Depending on the position of L3, this yielded effective FOV diameters anywhere between 2.5 and 3.5 mm. Importantly, in each case effective image brightness remained approximately constant across the full FOV (Fig. S1b, Methods). Here, the marginal brightness increase towards the edges is related to the slight upwards bend in the imaging plane as commonly seen for large FOV 2P microscopes (Sofroniew et al. 2016; Tsai et al. 2015). The axial difference between the edge and centre of the imaging plane was 20, 45, 87 and 170 μm for 1.2, 1.8, 2.5, 3.5 mm FOV, respectively. If required, this can be corrected via the ETL. However, biological structures are rarely perfectly flat either, so often a more useful solution might be to immediately fit the scan-plane to the 3D curvature of the interrogated sample (see below).

While these specific lens configurations readily work for the commercially available Sutter MOM, the fundamental concept of setting up an IFP to yield a diverging beam is readily applicable to any standard 2P microscope, provided the optical path between the scanning mirrors and tube lens is accessible. In fact, 2P scan-lenses tend to consist of multiple custom-designed optical elements which by themselves easily exceed the cost of our solution. Accordingly, if provided directly by the microscope’s manufacturer, our simplified optics should decrease the cost of such an off-the-shelf system.

### De-collimated optics facilitate rapid axial scans

In addition to expanding the FOV, our de-collimated design also shifts the excitation point beyond the objective’s nominal focal distance (Fig. S1c). Accordingly, the introduction of an electrically tunable lens (ETL) early in the laser path allows the user to exploit the same optical effect to drive rapid axial shifts in the excitation plane (Zhao et al. 2019). Specifically, we positioned an off-the-shelf ETL (EL-16-40-TC-20D, Optotune) 200 mm in front of the first scan mirror and controlled it with a custom driver board (see user manual). In this position, already a minor deviation from the perfectly flat curvature at zero input current slightly converged the laser which, in turn, strongly shifted the effective z-focus below the objective. For example, in both DBO_1_ and DBO_2_, stepping the input current from zero to 25% (50 mA) gave rise to a ~600 μm z-shift of the excitation plane. The use of only a small fraction of the ETL’s full dynamic range enabled short turnaround times (1-10 ms, depending on distance jumped, SFig. 3) and prevented overheating (Fahrbach et al. 2008; Grewe et al. 2011). If required, rapid synchronization of the ETL curvature with a Pockels cell for controlling effective laser power at the sample plane can compensate for any systematic variations in image brightness associated with increased penetration depth. Together, an investment of ~£1,000 (cf. user manual) therefore allowed expanding the standard Sutter MOM to a system capable of scanning anywhere within a 3.5×0.6 mm cylinder.

Our design’s full optical path and control logic are shown in Fig. 2d. All functions are executed from the scan software, which directly controls the xy-scan path as usual. To synchronize the ETL and/or a Pockels cell to this xy-scan, a copy of the fast-mirror command is sent to two microcontrollers. Each of these then executes preloaded line-synchronized commands that are defined using a standalone graphical user interface (GUI). In this way, this standalone z-control-system only requires a copy of the scan mirror command, meaning that it can be directly added to any 2P microscope setup without the need for software modifications.

### Divergent beam optics enable PSF-tailoring

To establish how our DBO approach affected the excitation PSF, we imaged 50 nm fluorescent beads across all configurations at 927 nm wavelength and constant laser power at the sample (Methods, Supplementary discussion). Starting from a DL spot-volume of 0.56 and 3.15 μm (xy and z, respectively), our different modifications elongated and laterally expanded the PSF laterally to varying degrees, from 0.77 and 9.94 μm for the 1.2 mm FOV configuration to 2.21 and 41.49 μm at 3.5 mm FOV (Fig. 2f-h). Notably, existing solutions for increasing the FOV also tend to present larger-than DL PSFs (Bumstead 2018; Sofroniew et al. 2016; Stirman et al. 2016; Tsai et al. 2015).

In DBO, increasing the FOV mainly elongated the PSF, while restricting its lateral expansion to remain comfortably suitable for providing single cell resolution in most biological samples even for the largest 3.5 mm expansion. FOV expansion also generally increased the system’s SBP (Fig. 2i).

The systematic effects on PSF shape also meant that our DBO approach could be used to flexibly tailor PSF dimensions to specific experimental needs. This can be achieved by varying the degree of underfilling of the objective’s back aperture while simultaneously keeping the laser’s divergence angle approximately constant. We demonstrate this principal capability by setting up a “high-resolution” (small PSF) variant of DBO_1_ (Fig. S2).

In general, such PSF-tailoring, is useful for balancing the spatial resolution with the SNR. For example, the sub-micron DL PSF offered by typical collimated 2P-setups maximises spatial resolution which is invaluable for resolving small synaptic processes or the somata of larval fruit flies (typically <5 μm). However, many species’ cell bodies and neural processes are much larger, as illustrated when imaging the brain of larval zebrafish, where a very small DL PSF spatially oversamples the “mid-sized” ~10 μm somata at the expense of a potentially substantial boost in SNR, a limitation that can be avoided by DBO-mediated tailoring of the PSF. Similarly, for picking up somatic signals from cortical neurons in the mouse, a “10fold expanded” ~5 μm PSF yields the best SNR (Weisenburger et al. 2019). Notwithstanding, any axial expansion in the PSF must be balanced with potentially undesirable merging of distinct image structures separated in depth.

As well as permitting the tailoring of the PSF to a given biological application, the use of a non-DL excitation spot can also bring about additional benefits. First, the lower effective excitation N.A. produces a narrower light cone which is less likely to be scattered by tissue inhomogeneities (Helmchen & Denk 2005). Second, objects that are smaller than the focal excitation volume become dimmer, while objects that are similar in size or larger remain bright (Birge 1986; Sofroniew et al. 2016). Third, PSF expansion also reduces photobleaching and photodamage which can have a more-than-quadratic intensity dependence (Hopt & Neher 2001; Patterson & Piston 2000). For example, when using the large PSF of the 3.5 mm FOV configuration, it was possible to use up to 250 mW laser power without causing notable damage when imaging deep in the mouse cortex (Podgorski & Ranganathan 2016).

### Increased effective laser power and working distance

Because our DBO design avoids overfilling of the objective’s back aperture and uses fewer optical elements in the laser path, total laser power at the sample was increased approximately 4-fold compared to all configurations of the DL setup (Fig. S1g). This additional power could, for example, be used to facilitate imaging deep in the brain, or alternatively to drive additional setups from the same laser source. For instance, when imaging the small brains of larval zebrafish or fruit flies, there is rarely a need to exceed 50 mW, meaning that it is theoretically possible to drive ten such DBO setups from a single standard laser (e.g. Coherent Chameleon Vision-S Laser, average power ~1.5 W at 930-960 nm, assuming 50% loss through the setup). Second, defocussing the laser also increased working distance below the objective, for example to provide additional space to position an electrode (Fig. S1c). However, this comes at the cost of slightly reducing photon capture by any detectors above the objective. Notably, photon capture efficiency from below the sample plane remains unaffected.

Taken together, our design therefore presents a low-cost and easily implemented solution to expand the FOV of any 2P microscope in three dimensions while maintaining image quality suitable for single cell resolution. In the following, we demonstrate how these capabilities can be exploited in a range of neurophysiological applications in mouse cortex, as well as the nervous systems of larval zebrafish and fruit flies.

## Part II. IMAGING THE STRUCTURE AND FUNCTION OF NEURONS

### High-resolution mesoscale and 3D random access imaging of the mouse brain

The width of the adult mouse’s brain is ~10 mm (Kovačević et al. 2005) which makes it too large to be comprehensively captured by conventional 2P microscopy. Here, an experimental goal might be to reliably resolve the ~20+ μm somata of major cortical or subcortical neurons across a 10 mm FOV. At the Nyquist detection limit, this would “only” require ~1,000 pixels across, which is well within the range of standard high-resolution scan-configurations.

Accordingly, currently the main limitation in achieving this goal is the microscope’s maximal FOV. Our DBO design makes important steps to address this limitation. When configured for a 3.5 mm FOV (DBO_2_), our setup captures about a third of the width of a mouse’s brain. In this configuration, a scan of a transverse section from a Thy1:GCaMP6f mouse (Fig. 3a,b) illustrates how the objective’s back aperture casts a shadow at the image edge, thus limiting the spatial extent of the scan (Fig. 3c). Within this maximal window, a high-resolution 1,024×1,024 px scan allowed us to resolve the somata of major cortical and hippocampal neurons (Fig. 1d, Supplementary Video S2). Accordingly, at this largest FOV configuration, effective signal detection largely suffices to capture the mouse brain’s major neuron populations. However, with our galvo-galvo setup, scan rates at this level of spatial detail were slow (0.49 Hz, 2 ms/line). Accordingly, we used a mesoscale imaging approach with reduced spatial sampling (256×256 px, 1 ms/line) to capture the entire image at 3.91 Hz. This permitted simultaneous population-level “brain-wide” recording of seizure-like activity across the cortex and underlying hippocampus following bath application of an epileptogenic (high K^+^, zero Mg^2+^) solution (Fig. 3e-g). To demonstrate the value of the system for more detailed readout of neuronal activity, we also used random access scans to simultaneously capture distant smaller scan-fields at high resolution, both spatially and temporally (two times 256×128 px at 3.91 Hz, Fig. 3d, h-j, Supplementary Video S2). In the example provided, the laser travelled between the two scan fields separated by ~1 mm within two 1 ms scan lines. This allowed us to record quasi-simultaneous neural activity across both the cortex and hippocampus at single cell resolution. The generally high SNR in these recordings also suggested that additional temporal or spatial resolution could be gained by the use of resonance scanners in place of our galvos (Fan et al. 1999).

The large FOV DBO_2_ configuration also lends itself to imaging mouse cortical neurons activity *in vivo* (Fig. 4), an increasingly common demand in neuroscience. Here, the maximal 3.5 mm FOV captured an entire cranial window of a Thy1-GCaMP6f mouse prepared for optical interrogation of the somatosensory cortex, comprising an estimated 10,000+ neurons in a given image plane (Fig. 4a-c, Supplementary Video S3, cf. Fig. 1g). Even in an intermediate DBO1 configuration (in this case a 1.5 mm FOV) the full image still comprised several 1,000s of neurons (Fig. 4d), many more than could be simultaneously captured at scan-rates suitable for functional circuit interrogation with a galvo-galvo setup. In an example scan we again used a random access approach to quasi-simultaneously record two 330 × 210 μm regions separated by ~1.2 mm (two times 128×64 px at 7.81 Hz). As in the brain slice preparation (Fig. 3), this reliably resolved individual neurons in spatially distinct regions of the mouse brain (Fig. 4d-g).

**Figure 4.**
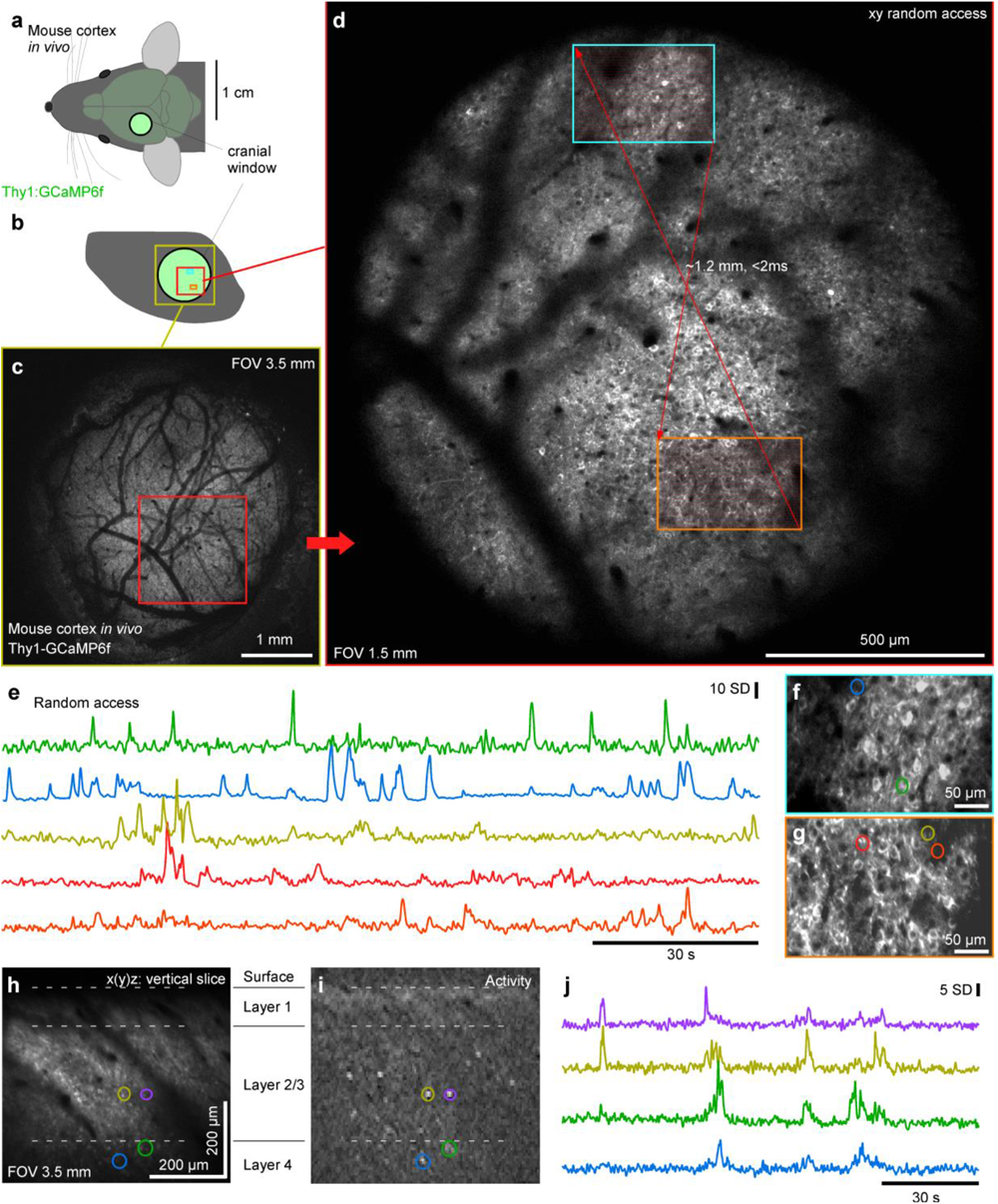
High-resolution mesoscale random access imaging: Mouse cortex in vivo. **a**,**b**, Schematic of Thy1:GCamP6f mouse brain in vivo (a) with cranial window over the somatosensory cortex (b). **c**,**d,** 1024×1024 px DBO_2_ (c) and DBO_1_ (d) images as indicated. **e-g**, 2 times 128×256 px (3.91 Hz) random access scan as indicated in (d) with fluorescence traces (e) as in (f,g). **h-j**, DOB1 128×128 px z-tilted plane (7.82 Hz) traversing through cortical layers 1-4 at ~27° relative to vertical with mean image (h), activity-correlation (i) and fluorescence traces (j) as indicated.

Finally, we also recruited the ETL to setup an axially tilted scan plane. This allowed quasi-simultaneous recording from neurons separated several hundreds of μm in depth across layers 1-4 of the mouse cortex *in vivo* (Fig. 4h-j).

### 3D plane-bending for imaging activity across the zebrafish brain

Owing to their small size and transparent larval stage, zebrafish have become a valuable model for interrogating brain-wide neural circuit function (Ahrens et al. 2013; Leung et al. 2013). However, at 7 *dpf* the brain is ~1.2 mm long and therefore too large to fit into the FOV of a typical DL 2P setup. Instead, its transparent body wall makes larval zebrafish well-suited for 1-photon selective-plane-illumination microscopy (1p-SPIM / “lightsheet microscopy”), which is not FOV-limited in the same way as 2P microscopy (Ahrens et al. 2013; Fahrbach et al. 2008; Sancataldo et al. 2019). However, 1p-SPIM and related techniques (Huisken & Stainier 2009) have a number of drawbacks, including constraints on However, during standard planar scans the powerful optical sectioning afforded by the 2P approach highlighted the 3D curvature of distinct brain regions by cutting right across them. While it was possible to image anywhere within the brain at high spatial resolution, the planar image grossly misrepresented the real 3D structure of the zebrafish brain (Fig. 5c, top panel). For example, the tectum in larval zebrafish is tilted upwards ~30° (Wulliman et al. 1996), meaning that rather than either cleanly sampling across its retinotopically organized surface, or perpendicularly across its stacked functional layers, the planar image instead cut the tectum at an effective 30° angle to yield a mixture of both, thus confounding interpretation. To ameliorate these issues, we used a 3D curved scan plane by driving the ETL as a sqrt(cosine) function of the slow y-mirror command (Methods). This enabled z-curvature “halfpipe” scans

**Figure 5.**
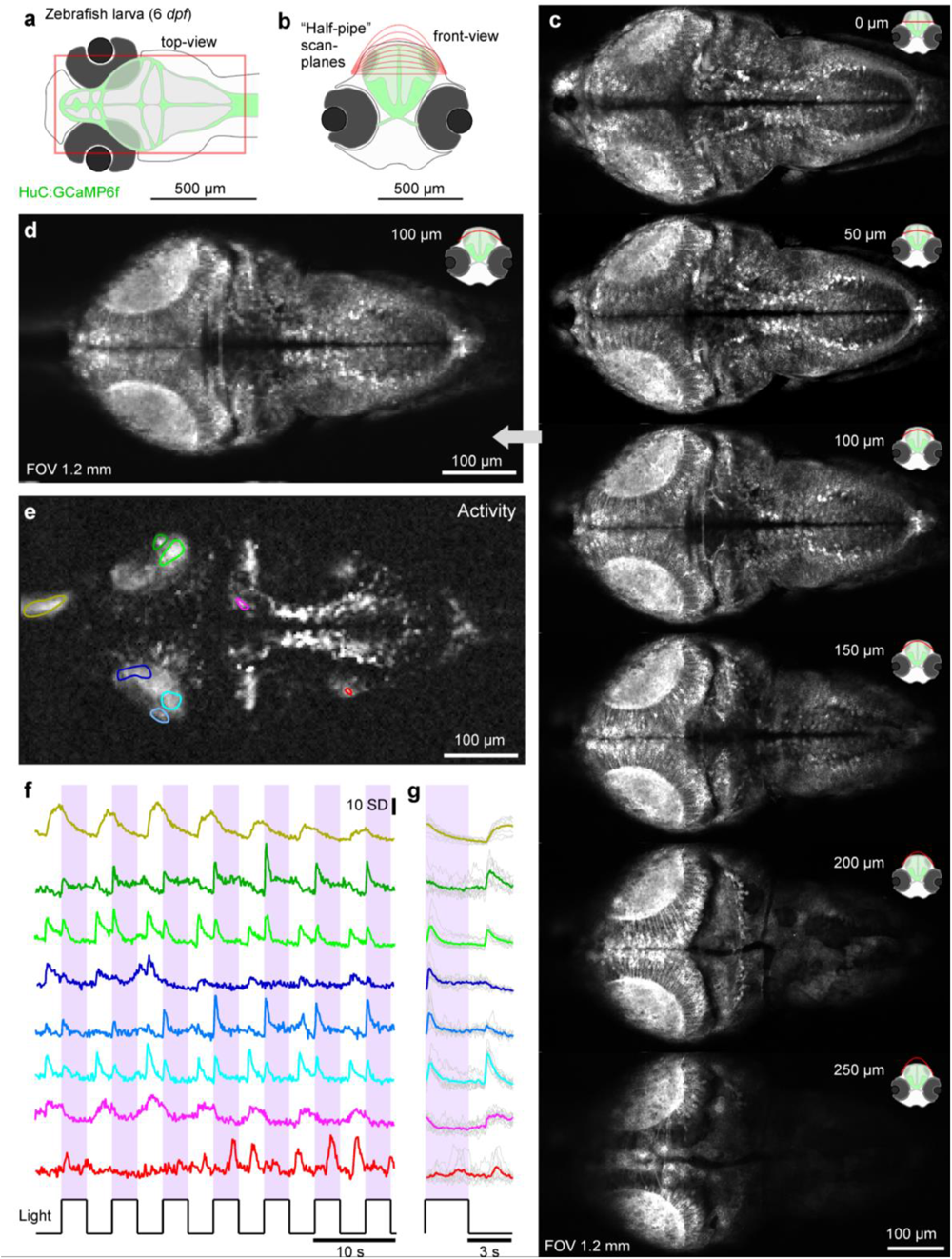
2P plane-bending to image the in vivo larval zebrafish brain. **a**, Schematic of HuC:GCaMP6f larval zebrafish brain viewed from top (a) and front (b) with scan planes indicated. **c**, DBO1 512×1024 scans of a 6 dpf zebrafish brain with different plane curvatures, with peak axial displacement at scan centre as indicated. At curvatures ~100-150 μm peak displacement the scan approximately traverses the surface of the tectum. **d-f**, mean (d), activity-correlation (e) and fluorescence traces (f, raw and g, event-triggered mean) from a 170×340 px scan (5.88 Hz) of the 100 μm peak displacement configuration (image 3 in (c)). The fish was presented with full-field and spectrally broad (~360–650 nm) series of light-flashes.

### 3D random access scanning across the zebrafish eye and brain

In the nervous achieving a homogenous image due to scattering and divergence of the excitation light with increasing lateral depth (Weisenburger & Vaziri 2018), limited access to tissues that are shadowed by strongly-scattering tissue such as the eyes (Hillman et al. 2019; Lavagnino et al. 2004) and, critically, a direct and bidirectional interference between the imaging system itself and any light stimuli applied for studying zebrafish vision (Vladimirov et al. 2014). These specific challenges could be readily addressed by our DBO 2P setup. For this, we used the smallest (1.2 mm FOV) configuration of DBO1, which just about fits one full zebrafish brain at a time while comfortably providing single cell resolution (Fig. 5a-c). that could be empirically fitted to follow the natural curvature of the brain, thereby closely capturing the functional anatomical organisation of the zebrafish brain (Fig. 5b,c, Supplementary Video S4). From here, we chose a single halfpipe plane that best followed the curvature of the two tecta and imaged this plane at 7.81 Hz (256×128 px, 1ms/line, Fig. 5d). We then presented spectrally-broad light stimulation that was synchronized to the scan retrace to prevent optical crosstalk. This allowed us to interrogate brain-wide visual function in response to arbitrary wavelength light (Fig. 5e-g). As required, the halfpipe scans could also be staggered for multiplane imaging at correspondingly lower image rates, including negative bends that surveyed the difficult-to-reach bottom of the brain between the eyes (Fig. S4, Supplementary Video S5). system, key functionally linked circuits are often separated in 3D space, representing a general problem for systems neuroscience. For example, the retinal ganglion cells of the zebrafish eye project to the contralateral tectum and pretectum, which are both axially and laterally displaced by several 100s of microns. Accordingly, it has been difficult to simultaneously record at both sites, for example to study how the output of the eye is linked to the visual input to the brain. To address this problem, we used our DBO1 configuration in synchronisation with the ETL to establish quasi-simultaneous 3D random access scanning of the zebrafish’s retinal ganglion cells across both the eye and brain (Fig. 6a,b).

**Figure 6.**
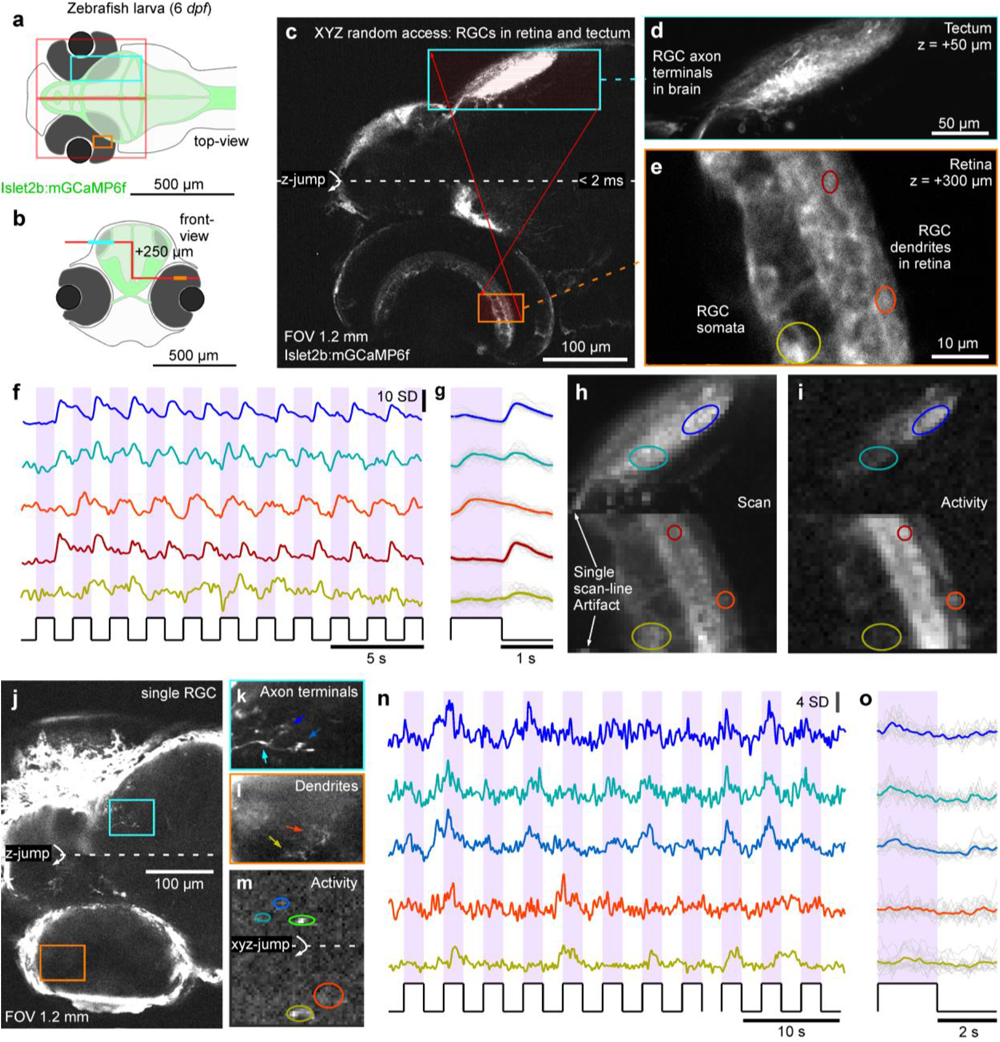
3D random access scanning of the zebrafish eye and brain. **a**,**b**, Schematic of zebrafish larva from top (a) and front (b) with scan configurations indicated. (c) DBO_1_ 1024×1024 px scan across an Islet2b:mGCaMP6f 6 dpf larval zebrafish eye and brain. At the centre of the scan, the axial focus is shifted by ~250 μm such that the axonal processes of retinal ganglion cells (RGCs) in the tectum (top) and their somata and dendritic processes in the eye (bottom) can be quasi-simultaneously captured. **d,e**, 1024×1024 px split-plane random access jump between tectum (d) and eye (e) and **f-i**, 2 times 64×128 px (15.6 Hz) random access scan of the same scan regions with raw (f) and event-averaged (g) fluorescence traces, mean image (h) and activity-correlation (i). The stimulus was a series of full-field broadband flashes of light as indicated. **j-o**, as (c-i), with individual RGCs transiently expressing GCaMP6f the same promoter.

For this we used an Islet2b:mGCaMP6f line which labels the majority of retinal ganglion cells in larval zebrafish. We first defined a slow, high-spatial resolution scan (512×512 px, 0.98 Hz) that captured the entire front of the head, however with a single z-jump at the centre of the frame to set-up a “staircase-shaped” scan-path (Fig. 6b,c). Here, empirical adjustment of the magnitude of the z-jump allowed us to identify the axonal processes of retinal ganglion cells in the brain, and their dendritic processes in the contralateral eye in the top and bottom of the same imaging frame, respectively. Based on this image, we next defined two scan regions for 3D random access scanning, one capturing a single plane across the tectum, while the other captured a smaller area of a subset of RGC dendrites and somata in the eye (Fig. 6c-e). Finally, we decreased the spatial resolution to 64×64 px to quasi-simultaneously image both regions at 7.81 Hz. This configuration allowed reliable recording of light-driven signals from individual RGC neurites across the eye and brain (Fig. 6f-i). Next, we repeated this experiment, however this time in zebrafish larvae that were transiently injected with Islet2b:mGCaMP6f plasmid. These animals stochastically express mGCaMP6f in only a very small number of RGCs, making it in principle possible in principle to identify the processes belonging to the same RGC in both the eye and brain. As a proof of principle, we present one such experiment where we could clearly image the processes of single RGCs at both sites (Fig. 6j-o). For this type of application, it will be important to optimise the genetic protocol to improve expression levels and thereby facilitate the identification of the same RGC’s processes at both sites.

### Multi-plane circuit mapping with optogenetics in Drosophila

Despite the generally enlarged FOV and concomitant increase in the *PSF*, our setup was still capable of resolving details of small neural processes in the <0.1 mm diameter nervous system of a first instar larval fruit fly. To assess the difference in image-resolution between our DBO setup and a DL-configuration, we first obtained anatomical scans from a third instar VGlut:GCaMP6f larva which expressed GCaMP6f in structurally well-defined neurons of the ventral nerve cord (Fig 7a,b). This revealed that while the DL image was clearly sharper (Fig. 7a), the DBO1 system nevertheless comfortably delineated individual somata (Fig. 7b). *Drosophila* was an ideal preparation to demonstrate our system’s capacity for multi-plane imaging for optogenetic functional circuit mapping (Fig. 7c-i). For this, we used a transgenic first instar larva that expressed the red-shifted optogenetic effector CsChrimson in all olfactory-sensory neurons (OSNs) on a background of pan-neuronal GCaMP6s (elav:GCaMP6s) (Fig. 7c,d). To reveal any potential bilateral crosstalk of olfactory signal processing across the brain’s two hemispheres, one of the olfactory nerves was cut. We set up six image planes (six times 340×170 px), each separated by ~20 μm which together captured the entire brain across both hemispheres at ~1 Hz (Fig. 7d, e). In this configuration, presentation of 2 s flashes of red light from a scanline-synchronized 590 nm LED activated olfactory sensory neurons (OSNs). These in turn propagated the signal to higher processing centres, which we visualised as regionally restricted GCaMP6s responses in the brain (Fig. 7f-I, Supplementary Video S6). The most strongly activated region was the ipsilateral antennal lobe (AL) which is directly innervated by the still-intact OSNs. Similarly, the olfactory second order processing centres, the mushroom body and the lateral horn, showed clear ipsilateral activation. In addition to these three major olfactory centres and their connecting tracts (e.g. plane 3), further processes and somata across both the ipsi-and contralateral lobe were also activated. Taken together, despite the slight expansion of the DL excitation spot, our DBO setup nevertheless allowed us to delineate key structural and functional information in this small insect brain.

**Figure 7.**
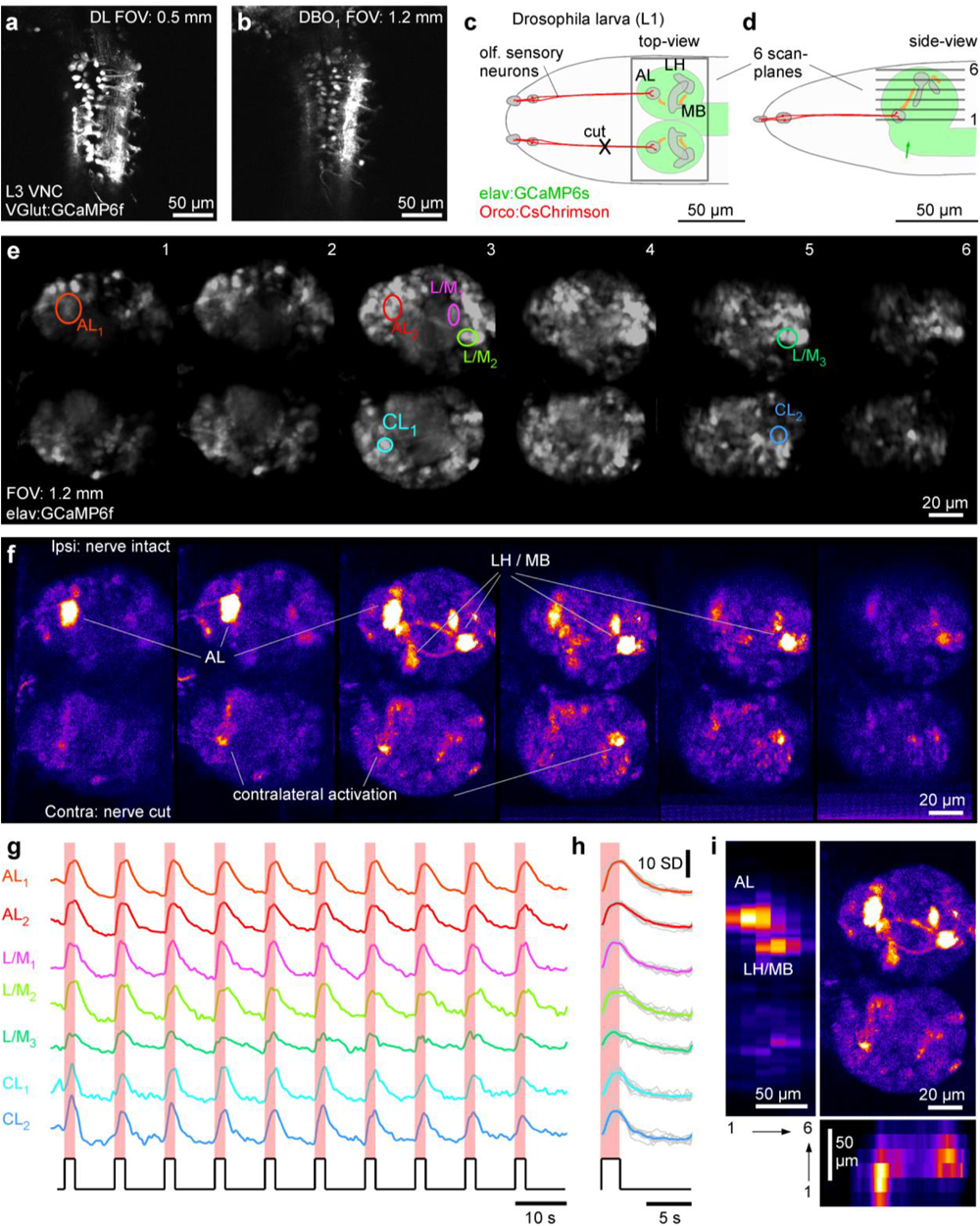
Multi-plane imaging and optogenetics for functional circuit mapping. **a**,**b**, DL (a) and DBO1 (b) 1024×1024 px scans of the ventral nerve cord of a 3^rd^ instar VGlut:GCamP6f Drosophila larva. **c**,**d**, Schematic of first instar elav:GCamP6s; Ocro:CsChrimson Drosophila larva from top (c) and side (d), with CsChrimson (red) and GCaMP6s (green) expression pattern as well as scan-planes indicated. **e-h** optogenetic circuit mapping of olfactory processing centres across the brain. Six scan planes (170×340 px each) were taken at 0.98 Hz/plane (i.e. volume rate) during presentation of 587 nm light flashes (2 s) to activate CsChrimson in olfactory sensory neurons (OSNs). Brain anatomy (e) and false-colour coded fluorescence difference image (f, 1-2 s after flash onset minus 1-2 s prior to flash onset), with fluorescence activity traces (g, raw and h, event triggered average). **i**, data from (f)summarised: top right: max-projection through the brain, with left and bottom showing transverse max-projections across the same data-stack.

## DISCUSSION

The ongoing development of sophisticated optical probes to report on key biophysical events has increasingly raised the demand in neuroscience for high SNR and large FOV 2P microscopes. To date, however, these characteristics are almost exclusively limited to high-end and, inevitably, high-cost platforms. Here, we exploit the fact that in 2P microscopy there is no “traditional” collection plane, allowing us to deviate from the diffraction limited regime that is typically used in systems where the planes of excitation and collection must superimpose to avoid image blur. Instead, we propose a simple core modification of the laser path that allows upgrading an out-of-the-box DL 2P microscope into a system capable of performing high SNR and large-FOV volumetric scans while at the same time preserving single cell resolution. We demonstrate the capabilities of this system for interrogating dynamic events in the brains of a range of key model species that are already widely used in neuroscience research. Since the core modification only requires the user to swap the scan lens for one or two off-the-shelf lenses, it can be tested (and fully reversed) within a matter of hours without the need for optical re-alignment or calibration. We anticipate that the simplicity and cost-effectiveness of this solution and the significant enhancement in 2P imaging capabilities that it permits, will lead to its wide adoption by the neuroscience community.

### Combining a DBO approach with existing custom 2P designs

The estimation of metrics that meaningfully compare the capability of our DBO design with other custom solutions is difficult, as these will generally depend strongly on the specific objective (N.A., back aperture size, working distance (focus)), its distance from the tube lens, and indeed the specifics of the interrogated sample and the biological question itself. Rather, because our DBO approach fundamentally differs from traditional DL optics, it opens the possibility to further enhance the capabilities of existing custom 2P microscope designs.

Moreover, our DBO approach is extremely flexible. It can be seamlessly implemented on setups with galvanometric or resonant mirrors to work with a wide range of scan-strategies. Here, the “extra” optical magnification afforded by the FOV expansion means that scan-mirror and ETL movements translate to relatively larger xy or z-translations, respectively, making it easy to rapidly execute complex and large-scale 3D scan-paths. Our approach can also be combined with existing setups that use rapid piezo-positioning of the objective for axial scans, although in this case the objective movement relative to the tube lens will generate small but systematic variations in FOV and PSF shape. Accordingly, the use of remote focussing before the scan-mirrors is likely to be preferable in most applications.

Like in most 2P designs, our use of a Gaussian beam does not permit the generation of a truly arbitrary PSF shape. Nevertheless, if used in combination with temporal focussing (Schrödel et al. 2013; Weisenburger et al. 2019) it would, in principle, be possible to modulate axial PSF expansion without strongly affecting lateral expansion, thus facilitating a greater range of PSF shapes. Similarly, an optimized design of the objective lens (Negrean & Mansvelder 2014) and other optical elements (Bumstead 2018) including the use of large diameter lenses to minimize aberrations (Tsai et al. 2015), could all be combined with our optical design to further enhance the quality of 2P excitation.

Taken together, our DBO approach offers key advantages over traditional DL 2P microscopy, including the capacity for an increased FOV, PSF-tailoring, rapid z-travel through minimal ETL commands and overall increased laser power at the sample plane. Moreover, it can principally be combined with a wide range of existing customisations to further push the capabilities of 2P microscopy in general. At the same time, our DBO approach is cost effective and can be readily implemented on an existing DL setup with minimal need for optical alignments and calibration.

## METHODS

### User manual

A complete user manual for the DBO design, as well as a bill of materials (BOM), 3D printable lens holders and printed circuit board (PCB) designs are available online at https://github.com/BadenLab/DBOscope.

### DL 2P microscope

Our setup was based on a Sutter MOM-type two-photon microscope (designed by W. Denk, MPI, Martinsried; purchased through Sutter Instruments) as described previously (Euler et al. 2009).

### Excitation path

The excitation beam was generated by a tuneable femtosecond Ti:Sapphire laser (Coherent Vision-S, 75 fs, 80 MHz, >2.5 W). The laser passed an achromatic half-wave plate (AHWP05M-980, Thorlabs) and was subsequently equally split to supply two independent 2P setups using a beam-splitter for ultrashort pulses (10RQ00UB.4, Newport). Next, the beam passed a Pockels cell (350-80 with model 302 driver, Conoptics), a telescope (AC254-075-B and AC254-150-B, Thorlabs), and was finally reflected into the head part of Sutter MOM stage by a set of three silver mirrors (PF10-03-P01). We used a pair of single-axis galvanometric scan mirrors (6215H, Cambridge Technology) which directed the beam into a 50 mm focal length scan lens (VISIR 1534SPR136, Leica) at a distance of 56.6 mm. A 200 mm focal length tube lens (MXA22018, Nikon) was positioned 250 mm further along the optical path. From here, the now collimated excitation beam was directed onto the xyz-movable head of the Eyecup scope (Euler et al. 2009) which was controlled by a motorized micromanipulator (MP285-3Z, Sutter Instruments). Here, the beam was reflected by two silver parabolic mirrors to pass the collection path dichroic mirror (T470/640rpc, Chroma) to finally slightly overfill the back aperture of the objective (Zeiss Objective W “Plan-Apochromat” 20x/1.0), thus creating a diffraction-limited excitation spot at the objective’s nominal working distance of 1.8 mm. The distance between the tube lens and the objective’s back aperture was 95 mm at the centre position of the xyz displacement mechanism, and the parabolic mirrors ensured that the optical excitation axes stayed aligned during movements of the microscope head.

### Collection path

Collection was exclusively through the objective. For this, a dichroic mirror (T470/640rpc, Chroma) was positioned 18 mm above the objective’s back aperture to reflect fluorescence light into the collection arm. Here, a 140 mm focal length collecting lens was followed by a 580-nm dichroic mirror (H 568 LPXR, superflat) to split the signal into two wavebands. The “green” and “red” channels each used a singleband bandpass filter (ET525/50 and ET 605/50, respectively, Chroma) and an aspheric condenser lens (G317703000, Linos) to focus light on a PMT detector chip (H10770PA-40, Hamamatsu).

### Image acquisition

We used custom-written software (ScanM, by M. Mueller, MPI, Martinsried and T. Euler, CIN, Tuebingen) running under IGOR pro 6.3 for Windows (Wavemetrics) to control the setup. For hardware-software communication we use two multifunction I/O devices (PCIe-6363 and PCI-6110, National Instrument). Within ScanM, we defined custom scan-configurations: 1,024×1,024 and 512×512 pixel images with 2 ms per line were used for high-resolution morphology scans, while faster, 1 ms or 2 ms linespeed image sequences with 256×256 (3.91Hz), 128×128 (7.81 Hz), 340×170 (5.88 Hz) or 128×64 (15.6 Hz) pixels were used for activity scans. All scans were unidirectional, and the laser was blanked via the Pockels cell during the turnarounds and retrace. This period was also used for light stimulation (zebrafish visual system and *Drosophila* optogenetics, see below).

### Non-collimated 2P microscope modifications

We used two sets of modifications (DBO1 and DBO_2_) to decollimate the excitation path to different degrees. For DBO_1_ (FOV 1.2 – 1.8 mm) we modified the original Sutter-MOM scan lens (VISIR 1534SPR136, Leica) by removing the second lens (i.e. the one closer to the tube lens) from the compound mount which changed the focal length from 50 to 190 mm. Alternatively, the entire de-constructed scan lens could also be replaced by a similar power off-the-shelf plano-convex lens. Our 190 mm lens (L1) was placed exactly 190 mm in front of the tube lens (so shifted 60 mm forward from its original position). Next, we introduced an additional plano-convex 175 mm focal distance lens (L2) (LA1229, Thorlabs). L2 was held in place by custom 3D printed mount (cf. user manual) inside the MOM’s tube-lens holder and positioned anywhere between 0 and 10 mm in front of the tube lens. Depending on the exact position of L2 within this range, the effective FOV at the image plane could be adjusted between 1.2 mm (10 mm distance) to 1.8 mm (L2 and tube lens almost touching).

For DBO_2_ (FOV 2.5 – 3.5 mm), we replaced the original scan lens with a single, 200 mm focal length plano-convex lens L3 (LA1708, Thorlabs). Like L2 in DBO_1_, L3 was mounted on the same custom 3D printed holder and positioned anywhere within a distance of 0-10 mm in front of the tube lens. In this case the FOV at the image plane could be adjusted between 2.5 mm (10 mm distance) to 3.5 mm (L3 and tube lens almost touching). For detailed instructions including photos of the optical path, consult the user manual.

We selected lens types and positions based on the available space within the Sutter MOM head such that for DBO1 and DBO_2_, the IFP was always located in front of or behind the TL, respectively. However, depending on the design of a given 2P setup’s excitation path, numerous alternative configurations are possible. Here, a straight-forward means to rapidly estimate the nature and scale of a given modification is to use a fluorescence test-slide and observe the change in working distance and FOV as the scan path is modified.

### Electrically tunable lens (ETL) for rapid axial focussing

For rapid z-focussing we added a horizontal ETL (EL-16-40-TC-20D, Optotune) into the vertical beam path after the silver mirror that reflected the excitation beam up into the MOM head, 200 mm in front of the scan-mirrors. To drive the ETL we used a custom current driver controlled by an Arduino Duo microcontroller (see user manual), capable of generating positive currents between 0-300 mA. The Arduino Duo received a copy of the scan-line command and in turn output commands to the current driver to effect line-synchronised changes in ETL curvature. Prior to initiating a scan, the specific to-be-executed Arduino programme was uploaded to the Arduino via serial from a PC running a custom Matlab-script (Mathworks). This Matlab script launched a simple graphical user interface (GUI) that allowed the user to configure the exact lens-path during a custom scan (see user manual). Accordingly, ETL control remained flexible and fully independent of the scan software. In this way, our solution can be readily integrated with any 2P system without need to change the software or acquisition/driver hardware. Notably, this ETL implementation can also be used by itself, without need for implementing any of the other optical adjustments described in this work. However, depending on the system’s optics, the effective range of z-travel would likely be smaller. A detailed step-by-step guide to implement the ETL, including the control software and hardware is provided in the user manual.

### Pockels cell

To control excitation laser intensity, we use a Pockels cell (Model 350–80, Conoptics; driver model 302, Conoptics). A line-synchronised blanking signal was sent from the DAQ to the drive to minimise laser power during the retrace. In addition, a custom circuit allowed controlling effective laser brightness during each scan line via a potentiometer (see user manual, designed by Ruediger Bernd, HIH, University of Tübingen). As required, this amplitude-modulated signal could then be further modulated by a second Arduino Due controlled by a standalone Matlab GUI to automatically vary effective laser power as a function of scanline index. In this way, laser power could be arbitrarily modulated on a line by line basis, for example to compensate for possible power loss when imaging at increased depth.

### Light stimulation

For visual stimulation of zebrafish larvae (Figs. 5, S2, 6) we used a full-field, broadband spot of light projected directly onto the eyes of the fish from the front via a liquid light guide (77555, Newport) connected to a custom collimated LED bank (Roithner LaserTechnik) with emission peak wavelengths between 650 and 390 nm to yield an approximately equal power spectrum over the zebrafish’s visual sensitivity range. LEDs were line-synchronised to the scanner retrace by an Arduino Due. For CsChrimson activation (Fig. 7) we used a custom 2P line synchronised LED stimulator (https://github.com/BadenLab/Tetra-Chromatic-Stimulator) equipped with four 587 nm peak emission LEDs embedded in a custom 3D printed recording chamber.

### Image brightness measurements

We imaged a uniform florescent sample consisting of two microscopy slides (S8902, Sigma-Aldrich) encapsulating a drop of low melting point agarose (Fisher Scientific, BP1360-100) mixed with low concentrated Acid Yellow 73 fluorescein solution (F6377 Sigma-Aldrich).

### PSF measurements

We used 175 ± 0.005 μm yellow-green (505/515) fluorescent beads (P7220, Invitrogen) embedded in a 1 mm depth block of 1% low melting point agarose (Fisher Scientific, BP1360-100). Image stacks were acquired across 30×30 μm lateral field of view with 256×256 pixels resolution (0.12 μm/pixel) and 0.5 μm axial steps/frame. For xy and z-dimensions, we calculated the full width at half maximum (FWHM) from Gaussian fits to the respective intensity profiles. Measurements were taken from set of the beads distributed across the entire FOV, and presented results are averages of at least 10 measurements of different beads, with error bars given in s.d..

### Animal experiments

All animal experiments presented in this work were carried out in accordance with the UK Animal (Scientific Procedures) Act 1986 and institutional regulations at the University of Sussex. All procedures were carried out in accordance with institutional, national (UK Home Office PPL70/8400 (mice), PPL/PE08A2AD2 (zebrafish)) and international (EU directive 2010/63/EU) regulations for the care and use of animals in research.

### Acute brain slices

1-2 month old male Thy1-GCaMP6f-GP5.17 (Dana et al. 2014) mice were used. Acute transverse brain slices (300 μm) were prepared using a vibroslicer (VT1200S, Leica Microsystems, Germany) in ice-cold artificial cerebrospinal fluid (ACSF) containing (in mM): 125 NaCl, 2.5 KCl, 25 glucose, 1.25 NaH_2_PO_4_, 26 NaHCO_3_, 1 MgCl_2_, 2 CaCl_2_ (bubbled with 95% O2 and 5% CO2, pH 7.3), and allowed to recover in the same buffer at 37°C for 60 minutes. During imaging, slices were constantly perfused with 37°C modified (epileptogenic) saline (37°C) containing 125 NaCl, 5 KCl, 25 glucose, 1.25 NaH_2_PO_4_, 26 NaHCO_3_, 2 CaCl_2_. Brain slices were imaged at 930 nm and 100-150 mW.

### Mouse surgical procedures for *in vivo*-imaging of the barrel cortex

Head bar implantation surgery has been described elsewhere (Bale et al. 2017). Briefly, under aseptic conditions, a male mouse expressing a calcium indicator in pyramidal neurons (GCaMP6f; GP5.17 (Dana et al. 2014)) was anaesthetised with isoflurane and implanted with a custom-made head bar. A circular 3 mm diameter craniotomy centred at 3.0 mm lateral and 1.0 mm posterior to bregma was made to expose the cranial surface. A cranial window, consisting of a 3 mm circular coverslip and a 5 mm circular coverslip (Harvard Apparatus), was placed over the craniotomy and secured in place with cyanoacrylate tissue sealant (Vetbond, 3M). Following 7 days of recovery, the mouse was handled daily and acclimated to a head fixation apparatus over a treadmill for a further 9 days. During 2P imaging, the head-fixed mouse could locomote freely on a custom-made treadmill. The mouse was awake and received fluid rewards between imaging batches. Cortical neurons were imaged at 960 nm and 100-150 mW.

### Zebrafish larvae preparation and *in-vivo* imaging

Zebrafish were housed under a standard 14:10 day/night rhythm and fed 3 times a day. Animals were grown in 200 mM 1-phenyl-2-thiourea (Sigma) from 1 day post fertilization (*dpf*) to prevent melanogenesis. Preparation and mounting of zebrafish larvae was carried out as described previously (Zimmermann et al. 2018). In brief, we used 6-7 *dpf* zebrafish (*Danio rerio*) larvae that were immobilised in 2% low melting point agarose (Fisher Scientific, Cat: BP1360-100), placed on the side on a glass coverslip and submerged in fish water. For eye-brain imaging, eye movements were prevented by injection of a-bungarotoxin (1 nL of 2 mg/ml; Tocris, Cat: 2133) into the ocular muscles behind the eye. Transgenic lines used were Islet2b:mGCaMP6f (eye-brain imaging) and HuC:GCaMP6f (Quirin et al. 2016) (image of 3 zebrafish in same FOV). Zebrafish were imaged at 930 nm and 30-60 or 50-100 mW for brain and eye imaging, respectively.

### Creation of Islet2b:mGCaMP6f transgenic line

*Tg(isl2b:nlsTrpR, tUAS:memGCaMP6f)* was generated by co-injecting pTol2-isl2b-hlsTrpR-pA and pBH-tUAS-memGaMP6f-pA plasmids into single-cell stage eggs. Injected fish were out-crossed with wild-type fish to screen for founders. Positive progenies were raised to establish transgenic lines. All plasmids were made using the Gateway system (ThermoFisher, 12538120) with combinations of entry and destination plasmids as follows: pTol2-isl2b-nlsTrpR-pA: pTol2pA(Kwan et al. 2007), p5E-isl2b(Pittman et al. 2008), pME-nlsTrpR(Suli et al. 2014), p3E-pA(Kwan et al. 2007); pBH-tUAS-memGaMP6f-pA: pBH(Yoshimatsu et al. 2016), p5E-tUAS(Suli et al. 2014), pME-memGCaMP6f, p3E-pA. Plasmid pME-memGCaMP6f was generated by inserting a polymerase chain reaction (PCR)-amplified membrane targeting sequence from GAP-43(Kay et al. 2004) into pME plasmid and subsequently inserting a PCR amplified GCaMP6f(Chen et al. 2013) at the 3’ end of the membrane targeting sequence.

### Drosophila larval preparation and in-vivo imaging

Flies were maintained at 25 °C in 12 h light:12 h dark conditions. Fly stocks were generated using standard procedures. The genotypes of the *D. melanogaster* flies used were: *elav-Gal4; LexAOp-CsChrimson* and *w*; *UAS-GCaMP6s; Orco-LexA*. These two strains were crossed to each other (collecting virgins from the first one and males from the second one) and placed on laying-pots at 25°C for larval collection. The laying-pots had a grape juice agar plate with an added drop of yeast paste supplemented with all-trans retinal (Sigma-Aldrich) to a final concentration of 0.2 mM. Yeast supplemented agar plates were changed every day and first instar larvae were picked off the new changed plate. First instar larvae were collected from yeast supplemented agar plates and dissected on physiological saline as in (Prieto-Godino et al. 2012) (in mM): 135 NaCl, 5 KCl, 5 CaCl_2_-2H_2_O, 4 MgCl_2_-6H_2_O, 5 TES (2-[[1,3-dihydroxy-2-(hydroxymethyl)propan-2-yl]amino]ethanesulfonic acid), 36 Sucrose, adjusted to pH 7.15 with NaOH. Larvae were dissected to expose the brain while maintaining intact the anterior part of the animal and the connection between OSN cell bodies and the brain, subsequently one of the olfactory nerves was cut with the forceps. The preparation was then positioned on top of a coverslip coated with poly-lysine (Sigma-Aldrich, P1524-100MG), and covered in 2% low melting point agarose (Fisher Scientific, Cat: BP1360-100) diluted in physiological saline, to prevent movement associated with mouth-hook contractions. The sample was then submerged in physiological saline. Larval brains were imaged at 930 nm and 30-60 mW.

## Supporting information

SVideo 1

SVideo 2

SVideo 3

SVideo 4

SVideo 5

SVideo 6

## Author contributions

FKJ and TB designed the study, with inputs from TE and all authors; FKJ implemented and tested hardware and software modifications, with input from PB, TY, TB and TE. FKJ and TB analysed the data, with inputs from all authors. PB assisted with hardware and software testing and troubleshooting and built the visual stimulator. MRB and MM provided mice for *in vivo* imaging and assisted with their handling and imaging. TY and MZ generated Islet2b:mGCaMP line and assisted with zebrafish sample preparation and testing. EK and KS provided mouse brain acute slice samples and assisted with handling and imaging. LLPG provided *Drosophila* sample and assisted with handing and imaging. TB built the optogenetics stimulator. FKJ and TB wrote the manuscript with inputs from all authors.

## Acknowledgements

We thank Sabi Abdul-Raouf Issa for providing the VGlut:GCaMP6f *Drosophila* sample, and John Bear for helping with the generation of the Islet2b:mGCaMP6f line. The authors would also like to acknowledge support from the FENS-Kavli Network of Excellence and the EMBO YIP.

## SUPPLEMENTARY MATERIALS

**Divergent excitation 2P microscopy for 3D random access mesoscale imaging at single cell resolution**, Janiak et al.

### A note on *PSF*-measurements on beads

Comparing the properties of 2P systems by way of describing their *PSFs* is complicated by the fact that the specific values obtained strongly depend on excitation power, wavelength, the specific size of the imaged fluorescent bead as well as their 2P excitation cross-section (Fig. S1d-f) (Göppert-Mayer 1931; Larson et al. 2003; Negrean & Mansvelder 2014; Zipfel et al. 2003). Moreover, one of the crucial parameters related to 2P absorption efficiency, the 2P cross-section, is for most fluorescent beads different from that of most popular biosensors for reporting neural activity (Ricard et al. 2018). Accordingly, *PSF*-measurements from fluorescent beads can only go part-way towards predicting the real effective excitation volume achieved in experiments with fluorescent biosensors.

**Figure S1, related to Figure 2.**
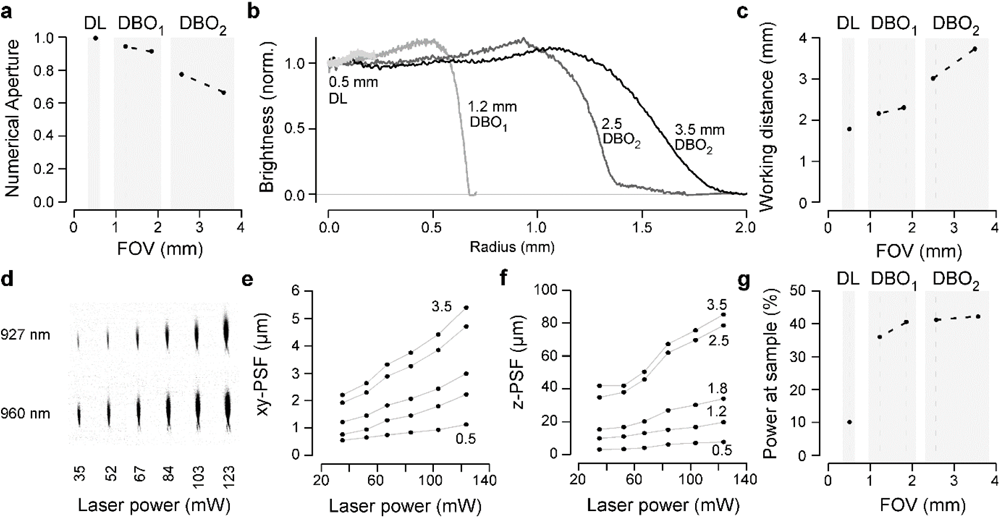
Quantification of DBO optical performance. Numerical Aperture (N.A.) calculated for the different optical configurations. **b**, Image brightness of the different configurations quantified from scans of a block of agarose with fluorescein. **c**, measured working distance. **d-g**, point-spread-functions (PSF) measurements at varying laser power and wavelength as indicated. **g**, Power at sample measured for all configurations, expressed as a percentage of the power that reaches the scanning mirrors.

**Figure S2 | related to Figure 2.**
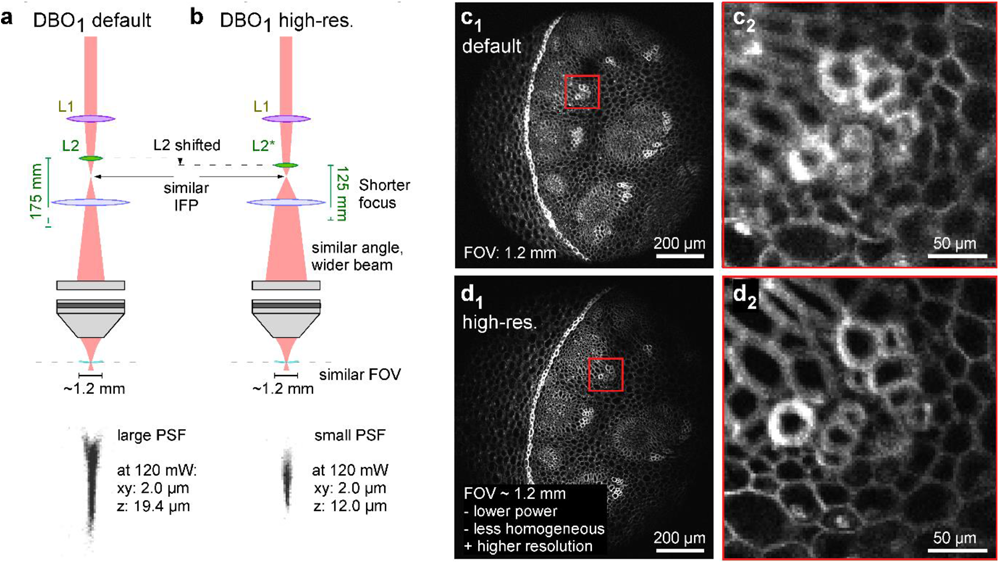
Tailoring PSF independent of FOV. **a**, default DBO_1_ configuration and **b**, optical modification to achieve a smaller PSF (bottom) while maintaining a ~1.2 mm FOV. To achieve this effect, L2 (175 mm FL) was exchanged for L2*, (125 mm FL). L2* was also shifted closer towards the TL to set up a similar IFP compared to the default configuration. As a result, a now expanded laser beam (hence smaller PSF) reaches the objective’s back aperture at a similar divergence angle (e similar FOV). **c,d**, Fluorescence test slide imaged under either configuration at 1,024×1,024 px resolution and full FOV (c_1_,d_1_), and inset enlarged as indicated (c_2_,d_2_).

**Figure S3 | related to Figure 2.**
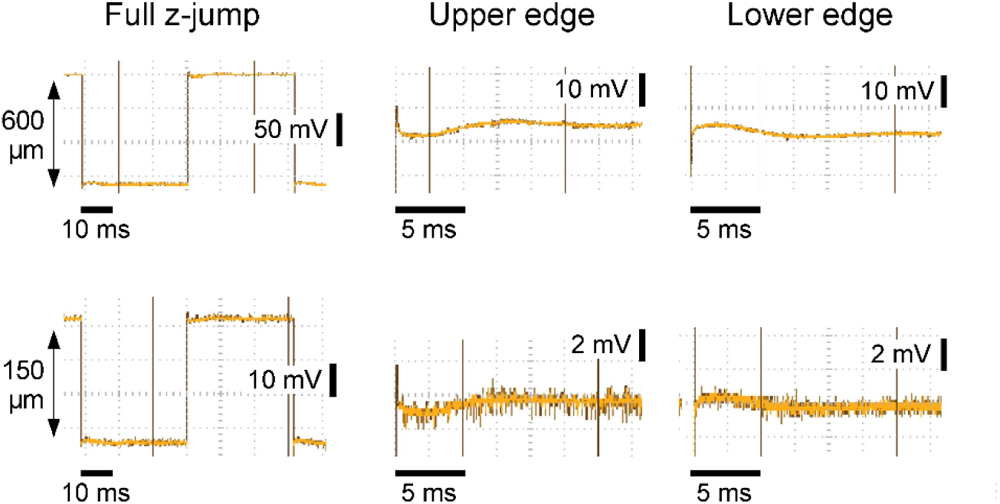
ETL settling time. Voltage on the electrically tunable lens (ETL) recorded on an oscilloscope through a resistor as a readout of current curvature. In response to current step commands that resulted in 600 (top) and 150 (bottom) μm axial focus jumps, the lens oscillated with <5% maximal jump amplitude following an initial sub-millisecond transient. This oscillation reliably settled beyond detection limit within <10 ms (600 μm jump). For smaller jumps it settled substantially earlier.

**Figure S4 I related to Figure 5.**
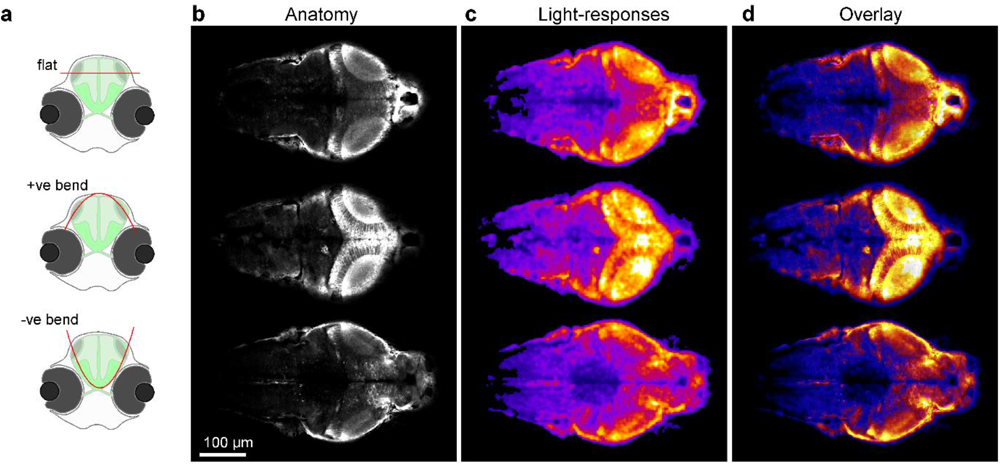
Staggered plane bending. **a**, Schematic of HuC:GCaMP6f larval zebrafish shown from front, with scan-planes indicated. b-d, three times 170×340 px (1.96 Hz volume rate) staggered bent-planes used to quasi-simultaneously capture the brain at three different orientations as indicated, with mean image (b), fluorescence-difference image (c, 1-2 s after stimulation light onset minus 1-2 s prior to stimulation light onset, with stimulus as in Fig. 5) and overlay (d).

## SUPPLEMENTARY VIDEOS

**Supplementary Video S1, related to Figure 1 | Z-stack through three larval zebrafish**. DBO_2_ z-stack (3.5 mm FOV) of three larval zebrafish as in Fig. 1f_1_, 1,024×1,024 px (0.5 Hz per plane), 1 μm z-steps.

**Supplementary Video S2, related to Figure 3 | Mesoscale imaging of mouse brain slice**. DBO_2_ scan (3.5 mm FOV) of seizure-like activity mouse brain slice, in 2 parts. First, FOV as shown in Fig. 3d (1,024×1,024 px, 0.5 Hz), and second as shown in Fig. 3h,i. (2 times 128×256 px, 3.91 Hz each).

**Supplementary Video S3, related to Figure 4 | Mesoscale imaging of mouse cortex *in vivo***. As in Fig. 4d, DBO_1_ scan (1.5 mm FOV) of spontaneous activity in mouse somatosensory cortex at 1,024×1,024 px (0.5 Hz).

**Supplementary Video S4, related to Figure 5 | Half-pipe imaging of larval zebrafish brain**. As in Fig. 5c, DBO_1_ anatomical scan (1.2 mm FOV) of larval zebrafish brain, with increasing z-curvatures applied. 512×1,024 px (1 Hz per plane). Note that planes 3 and 4 most closely follow natural brain curvature.

**Supplementary Video S5, related to Figure S4 | Half-pipe multiplane imaging of larval zebrafish brain**. As in Fig. S4, DBO_1_ scan (1.2 mm FOV) of larval zebrafish brain, with three different z-curvatures applied: none (top) positive (middle) and negative (bottom). 512×1,024 px (1 Hz per plane).

**Supplementary Video S6, related to Figure 7 | Multiplane optogenetics in *Drosophila* larva**. As in Fig. 7e,f, DBO_1_ scan (1.2 mm FOV) of L1 larval *Drosophila* brain, with six planes scanned at 170×340 px, 0.98 Hz volume rate during optogenetic activation of CsChrimson in olfactory sensory neurons. Average of 10 stimulus repeats. (left: fluorescence average, right, background subtracted and false colour-coded).

